# Herbal medicine of Kursi Wufarikun Ziyabit inhibits mitochondrial ATP production to activate AMPK in hepatocytes in therapy of type 2 diabetes

**DOI:** 10.1101/2020.09.28.316349

**Authors:** Mireguli Yasen, Lan Jiang, Mizhgona Sharofova, Xuelei Xin, Yusuf Nuraliev, Jianping Ye, Haji Akber Aisa

## Abstract

The herbal medicine of Kursi Wufarikun Ziyabit contains two herbs (*Geranium collinum* Steph.ex Willd and *Hypericum Scabrum* Lnn) in the formula of ethnomedicine. It is usually used in the treatment of diabetes mellitus with a significant therapeutic efficacy. However, the molecular mechanism remains unknown for the action of medicine. In this study, we investigated the mechanism with a focus on an extract (GC30) of the medicine in the regulation of energy metabolism in hepatocytes. GC30 was prepared from a crude extract of the medicine through elution of an AB-8 macro porous resin column loaded with the medicine in 30% ethanol. GC30 exhibited an activity in the inhibition of triglyceride (TG) accumulation in the mouse hepatocytes through a suppression of SREBP1c activity. FGF21 (fibroblast growth factor 21) expression was induced by GC30 in a dosage-dependent manner at concentrations of 25 - 100 μg/ml. The induction was observed in mRNA and protein of FGF21, which were peaked at 2 hours and lasted for 8 hours in the response to GC30 (100 μg/ml). The transcription of FGF21 gene was induced by GC30 for an increase in the FGF21 gene promoter activity. AMPK and PKA activities were induced by GC30 with an elevation in their phosphorylation status, which were associated with a reduction in ATP abundance and an increase in CREB phosphorylation in cells treated with GC30. Oxygen consumption of mitochondria was inhibited in the hepatocytes by GC30. These activities of GC30 were similar to those of diabetes medicines including metformin and berberine. The data suggest that GC30 inhibited ATP production in mitochondria to activate AMPK and PKA in the hepatocytes to induce FGF21 expression. This study suggests a novel activity of the herbal medicine in the regulation of glucose and lipid metabolism in the hepatocytes.

## Introduction

Herbal medicines represent a resource of natural compounds in the control of blood glucose in the treatment of diabetes ^[1]^. Folk medicines provide thousands of drug candidates with activities in the control of diabetes. In this case, a formula herbal medicine by the name of Kursi Wufarikun Ziyabit is an ethnomedicine from Avicenna’s “Canon of Medicine”, which has a long history in the treatment of diabetes mellitus. The formula contains two different herbs, *Geranium collinum* Steph.ex Willd and *Hypericum Scabrum* Lnn. They are common herbs in the countries of Central Asia. *Geranium collinum* is a perennial plant that belongs to the family of Geraniaceae, which grows in the tropical and subtropical areas with 810 species in 8 genera in the world. Decoction of the *Geranium collinum* roots has a record in the treatment of the gastric diseases, postpartum bleeding, and hemoptysis as an astringent. In China, Tajikistan and Mongolia, various types of geranium are used in the treatment of hepatic and gastrointestinal diseases in the folk medicines. The water extracts of *Geranium collinum* roots are also used in the treatment of inflammation-related diseases, such as rheumatism and gout. Aqueous extract of the aerial part of *Geranium collinum* (from the flowers and leaves) are widely used in the treatment of eye diseases in Tibetan and Mongolian traditional medicine. The alcohol extract of the aerial part of *Geranium collinum* has a potent anti-inflammatory effect in the treatment of pneumonia, catarrhal symptoms of stomach and intestines. *Geranium collinum* contains phenols (pyrogallol, catechol), saponins, tannins, carbohydrates (monosaccharides, sucrose, glucose, fructose, sorbose, rhamnose, arabinose, xylose, ribose, maltose, starch, hemicellulose), vitamin C, flavonoids (avikulyarin, quercetin, gvayaverin, kaempferol, etc.). The molecular basis of the Kursi Wufarikun Ziyabit activity remains unknown in the treatment of diabetes mellitus. In this study, we addressed this issue by testing the effect of Kursi Wufarikun Ziyabit on hepatocytes.

AMP-activated kinase (AMPK) is a molecular target in the control of blood glucose in the type 2 diabetes ^[2]^. AMPK is activated in the liver by the diabetic medicines, such as metformin and berberine, to reduce the fasting blood glucose. The mechanism is related to inhibition of gluconeogenesis in hepatocytes to decrease glucose production by the liver ^[3]^. AMPK suppresses transcription-dependent expression of glucogenic genes by targeting the transcription factors (FOXO1 and CREB, etc.) ^[2]^. AMPK inhibits de novo lipogenesis through a similar mechanism by targeting expression or phosphorylation of lipogenic proteins (SREBP and ACC, etc.) ^[2]^. A reduction in AMPK activity leads to production of glucose by liver, which represents a mechanism for the hyperglycemia in type 2 diabetes. Activation of AMPK is a molecular mechanism for the therapeutic activities of metformin and berberine in the treatment of type 2 diabetes ^[4]^. However, the activity of Kursi Wufarikun Ziyabit remains unknown in the regulation of AMPK.

FGF21 is a fasting-induced hepatokine that promotes energy expenditure through an action in multiple peripheral tissues including adipose tissue ^[5, 6]^. Blood FGF21 is predominantly secreted by the liver although other tissues including adipose tissue, pancreas, and skeletal muscle also produce FGF21 ^[7, 8]^. FGF21 induces beige cell functions through activation the transcription factor PGC-1 ^[5]^. The FGF21 action is mediated by a FGF receptor complex containing β-klotho. Administration of recombinant FGF21 or transgenic overexpression of FGF21 in mice provides a protection against diet-induced obesity and glucose disorders ^[9]^. The activity of Kursi Wufarikun Ziyabit is unknown in the regulation of FGF21 expression.

In this study, we report that GC30 fraction of Kursi Wufarikun Ziyabit inhibited hepatic steatosis and induced FGF21 expression in hepatocytes. The mechanism was associated with an activation of AMPK through inhibition of ATP production in mitochondria. The study provides a cellular and molecular mechanism for the pharmacological activities of Kursi Wufarikun Ziyabit in the treatment of diabetes.

## Methods

### Plant material

The two herbs *Geranium collinum* Steph and *Hypericum scabrum* Lnn. were collected from the Botanical Garden, Academy of Sciences of the Republic of Tajikistan (38.5357500 N, 68.7790500 E and 767 m above sea level, Dushanbe, Tajikistan) in June 2016. The plants were identified by Qurbonov Mansur. A voucher specimen has been deposited in Xinjiang Technical Institute of Physics and Chemistry, Urumqi, Chinese Academy of Sciences.

### Preparation of GC30 extract

The medicine of Kursi Wufarikun Ziyabit was formed with *Geranium collinum* Steph (35g) and *Hypericum scabrum* Lnn. (15g) in the formula of this study. The herbs were grinded into powders individually and mixed at the ratio 2.3:1 as a powder mixture. The mixture was extracted with 50% ethanol at 70 □. The GC30 extract was obtained from a 50% ethanol extract of the powder mixture following a purification process on an AB-8 macro porous resin column, which was washed successively with distilled water (1BV), 30% ethanol (2BV), and 70% ethanol (3BV). The eluents of 30% and 70% ethanol were concentrated under reduced pressure and named as named GC30 and GC70, respectively. The compounds in GC30 were determined as described in supplementary material.

### Cell culture and reagents

The mouse hepatocyte cell line Hepa-1c1c7 (1c1c7 in this study) and HepG2 cells were purchased from the American Type Culture Collection (ATCC) (Manassas, VA 20110) and maintained in Dulbecco’s Modified Eagle’s Medium (DMEM) supplemented with 10% fetal bovine serum at 37 °C in a 5% CO_2_ incubator. Hepatic steatosis was established in HepG2 cells with a treatment by oleic acid (200μM) for 72 hr. The cells were treated with GC30 after serum-starvation in DMEM medium containing 0.25% fatty acid-free bovine serum albumin overnight. Berberine and metformin were used as positive controls in this study. Berberine (BBR, 200275,), Metformin (Met, PHR1084, Sigma-Aldrich), Geraniin (GN, E-0742, Tauto Biotech) and Ellagic acid (EA, E-0654, Tauto Biotech) were purchased from the corresponding commercial sources.

### Western blot

FGF21 Protein and AMPK phosphorylation were determined in Western blot as described elsewhere ^[10]^. The whole cell lysate was prepared in a lysis buffer with sonication as described elsewhere ^[10]^. Antibodies to FGF21 (ab171941) and actin (ab6276) were from Abcam (Cambridge, MA). Antibodies to SREBP1c (sc-8984, Santa Cruz Biotech., USA), pAMPKα (Thr172, #2531, Cell Signaling Biotech., USA) and AMPKα (sc-398861, Santa Cruz Biotech., USA) were purchased from the commercial sources.

### mRNA expression

FGF21 mRNA was determined in cells using qRT-PCR. Total RNA was prepared from the cells using TRIzol reagent following the manufacturer’s protocol (Invitrogen, Carlsbad, CA). *Fgf21* mRNA was determined in mouse 1c1c7 cells with the SYBR green primer (forward 5a-CCTCTAGGTTTCTTTGCCAACAG-3; reverse 5-AAGCTGCAGGCCTCAGGAT-3) ^[11]^. Target mRNA was normalized to ribosome 18S RNA in qRT-PCR. SYBR Green Master Mix (4309155, Applied Biosystems) was used in qRT-PCR with the ABI 7900 machine.

### Lipids assay

Cells were lysed in PBS (1 g: 20 mL). The lipids were extracted from the lysate using a chloroform/methanol (2:1) mixture ^[12]^. TG and glycerol were determined using the Serum Triglyceride Determination Kit (TR0100; Sigma-Aldrich). Oil red O staining was performed in HepG2 cells using a protocol described elsewhere ^[13]^.

### Mitochondria and ATP assays

Mitochondrial activity was determined in response to GC30 treatment by monitoring the oxygen consumption rate (OCR) using a XF24 Analyzer (Seahorse Bioscience Inc., North Billerica, MA). OCR was determined in the cells following 1 h pretreatment by GC30 in the serum-free DMEM medium. The assay was conducted in the XF base medium (Cat # 102353-100, Agilent Seahorse XF) supplemented with glucose (25 mM), pyruvate (1 mM) and glutamine (4 mM) to fuel the cells. In the control, berberine (10 μM) was used as a positive control for mitochondrial inhibitor. Following chemicals were used at a final concentration in the assay to modulate the respiratory chain: oxidative phosphorylation uncoupler (Carbonyl cyanide 4-trifluoromethoxy phenylhydrazone, FCCP, 2 μM), the complex V inhibitor (Oligomycin, 1 μM), complex II and III inhibitor (Antimycin A, 0.5 μM) and complex I inhibitor (Rotenone 0.5 μM). ATP production by mitochondria was determined in two ways. The first was a luciferin-based ATP assay (ATP Determination Kit, Life technologies, Eugene, Oregon, USA) as described elsewhere ^[14]^ and the second was determined by the value of OCR change following GC30 treatment.

### FGF21 gene reporter assay

A FGF21 luciferase reporter (0.2 mg DNA per well) was transfected into HepG2 cells in a 24-well plate (4 × 10^5^ cells per well) using Lipofectamine 2000 (Cat #11668027, ThermoFisher Scientific, Waltham, MA, USA). In the reporter vector, the luciferase gene was under control of a mouse FGF21 gene promoter (−1497/+5) as described elsewhere ^[15]^. The reporter activity was determined using the dual luciferase substrate system (E1910; Promega, Madison, WI, USA), and normalized with the Renilla luciferase. Each experiment was repeated three times with consistent results.

### Statistical analysis

In this study, the data were presented as the mean ± SEM. The experiments were repeated at least three times with consistent results. The representative immunoblotting is presented with quantification of three blots. Student’s T test was used in the data analysis with significance at *p* <0.05.

## Results

### Inhibition of TG accumulation in hepatocytes by GC30

To study the metabolic activity of herbal medicine of Kursi Wufarikun Ziyabit, GC30 was extracted from the powder of herb medicine and tested in hepatocytes. In the study, a hepatic steatosis model was established in the mouse 1c1c7 hepatocyte cell line by treatment of the cells with oleic acid for 72 hr (OA, 200 μM) (Fig. 1A). The TG abundance was quantified in the hepatocytes using the oil red O staining and a TG assay kit. In this model, treatment of the cells with GC30 during the induction of steatosis led to a significant reduction in the TG content, and the activity was comparable to the berberine activity (Fig. 1, A-C). Berberine was used as a positive control in the inhibition of TG accumulation ^[16]^. GC30 was used at 50 μg/ml in this study. When GC30 was applied to the cells after establishment of hepatic steatosis, the GC30 activity was not observed in the reduction of TG content (Fig. 1D), suggesting that GC30 inhibits lipogenesis and does not induce lipolysis. BBR also failed to reduce TG in the established hepatic steatosis (Fig. 1D). These data suggest that GC30 inhibits TG accumulation in hepatocytes, and the mechanism is related to the inhibition of TG synthesis. The GC30 activity is comparable to that of BBR.

**Fig 1.**
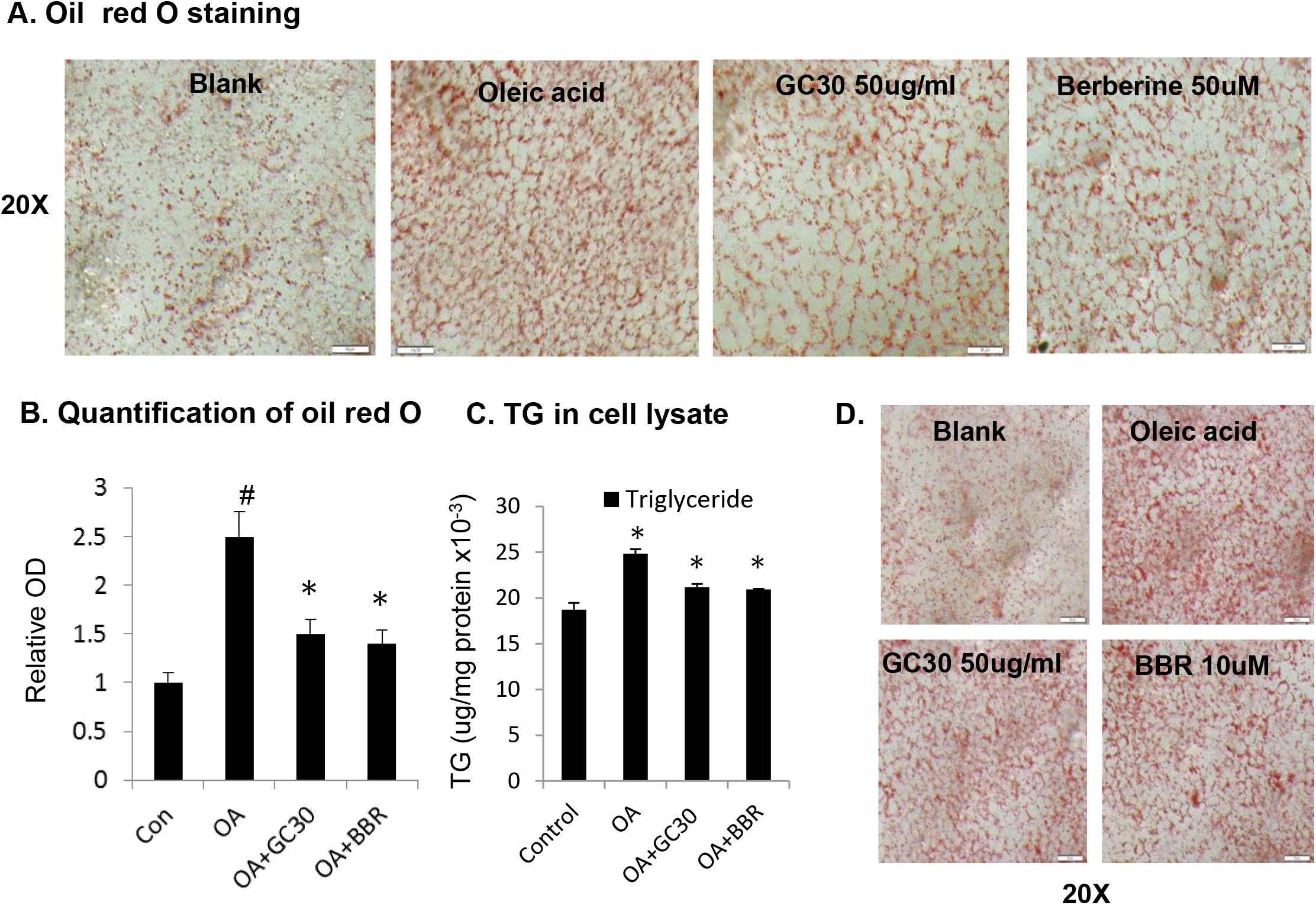
Inhibition of TG accumulation by GC30 in HepG2 cells. HepG2 cells were treated with oleic acid (200 μM) for 72 hours to induce TG accumulation in the hepatic steatosis model. GC30 was used at 50 μg/ml in this study. A. TG detection with oil red O staining. Lipid droplets are indicated by the red color in the HepG2 cell cytoplasm following oil red O staining. GC30 was applied to the cells together with the oleic acid during induction of TG accumulation. B. TG quantification by OD of color change. OD was determined in the extract of hepatocytes for oil red O. C. TG quantification by the TG assay kit. TG was extracted and quantified with the TG assay kit. D. GC30 treatment of established steatosis. In the bar figure, the results are mean ± SE (n=3). ^*^ p < 0.05; ^#^ p < 0.05 over the control.

### Inhibition of SREBP1c protein by GC30

In the hepatocytes, the transcription factor SREBP1c is one of the major driving forces in lipogenesis of hepatic steatosis. SREBP1c induces expression of multiple lipogenic enzymes through gene transcription in the induction of hepatic steatosis. SREBP1c protein was examined in the GC30-treated cells by Weston blotting to understand the mechanism of lipogenesis inhibition. In a 24 hr study, SREBP1c was decreased in the cells by GC30 in a time-dependent manner (Fig. 2). A significant reduction was observed after 16 hr of treatment. In the control, BBR also reduced SREBP1c protein in the model (Fig. 2). The data suggest that GC30 inhibits SREBP1c activity in hepatocytes to suppress lipogenesis in the hepatic steatosis model.

**Fig 2.**
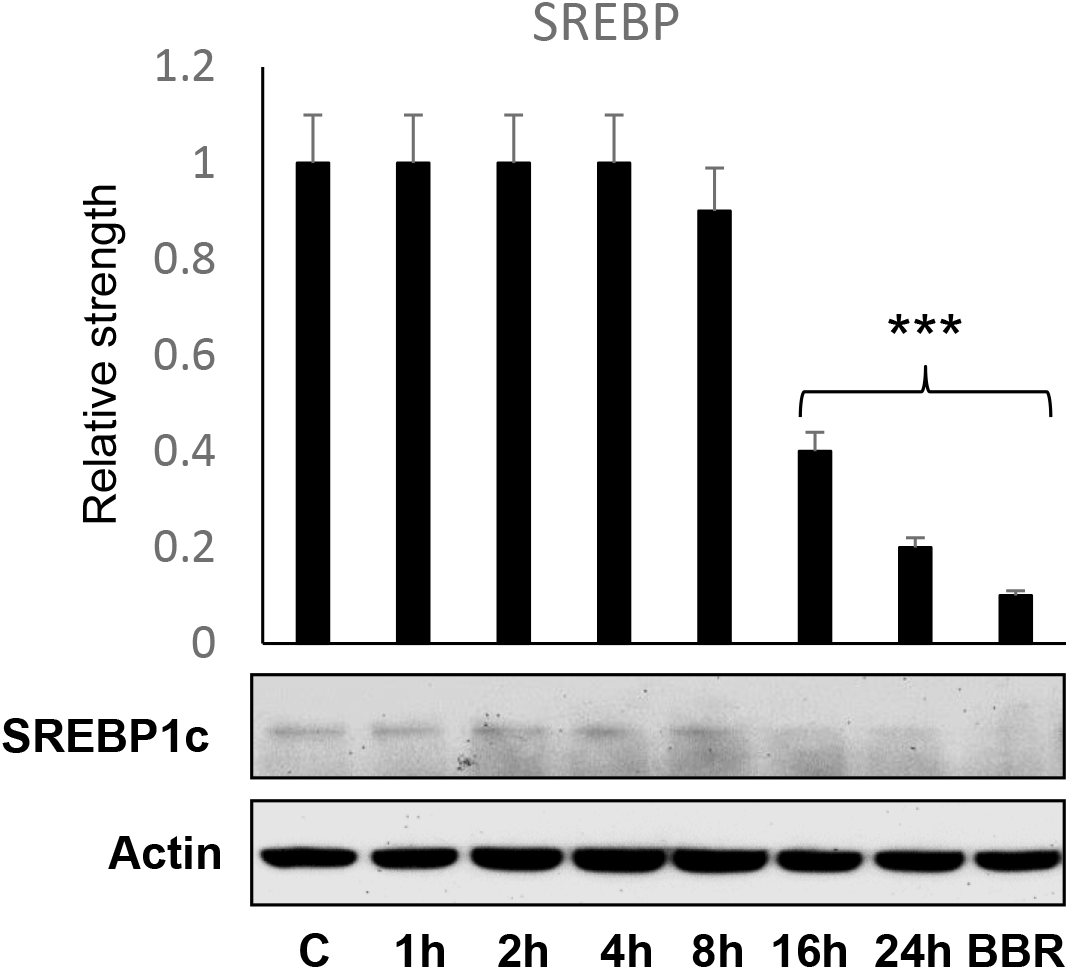
Inhibition of SREBP1c protein expression in hepatocytes by GC30. SREBP1c protein was examined in HepG2 hepatocytes after GC30 (100 μg/ml) or BBR (10 μM) treatment by Western blot. The bar figure represents mean ± SE (n=3). ^***^ p < 0.001 over the control.

### Induction of FGF21 expression by GC30

FGF21 is a major endocrine factor from liver with an activity in the regulation of whole-body energy metabolism. FGF21 was examined in this study to determine GC30 activity in the regulation of endocrine function of hepatocytes. FGF21 protein was examined in the cell lysate by Western blotting. The protein was induced by GC30 in a dose-dependent manner and the highest activity was observed at dosages of 50 - 100 μg/ml (Fig. 3A). In the time-dependent study, FGF21 was induced by GC30 during 1 - 24 hr. The induction was peaked at 4 hr (Fig. 3B). These results suggest that GC30 may induce FGF21 expression in hepatocytes.

**Fig 3.**
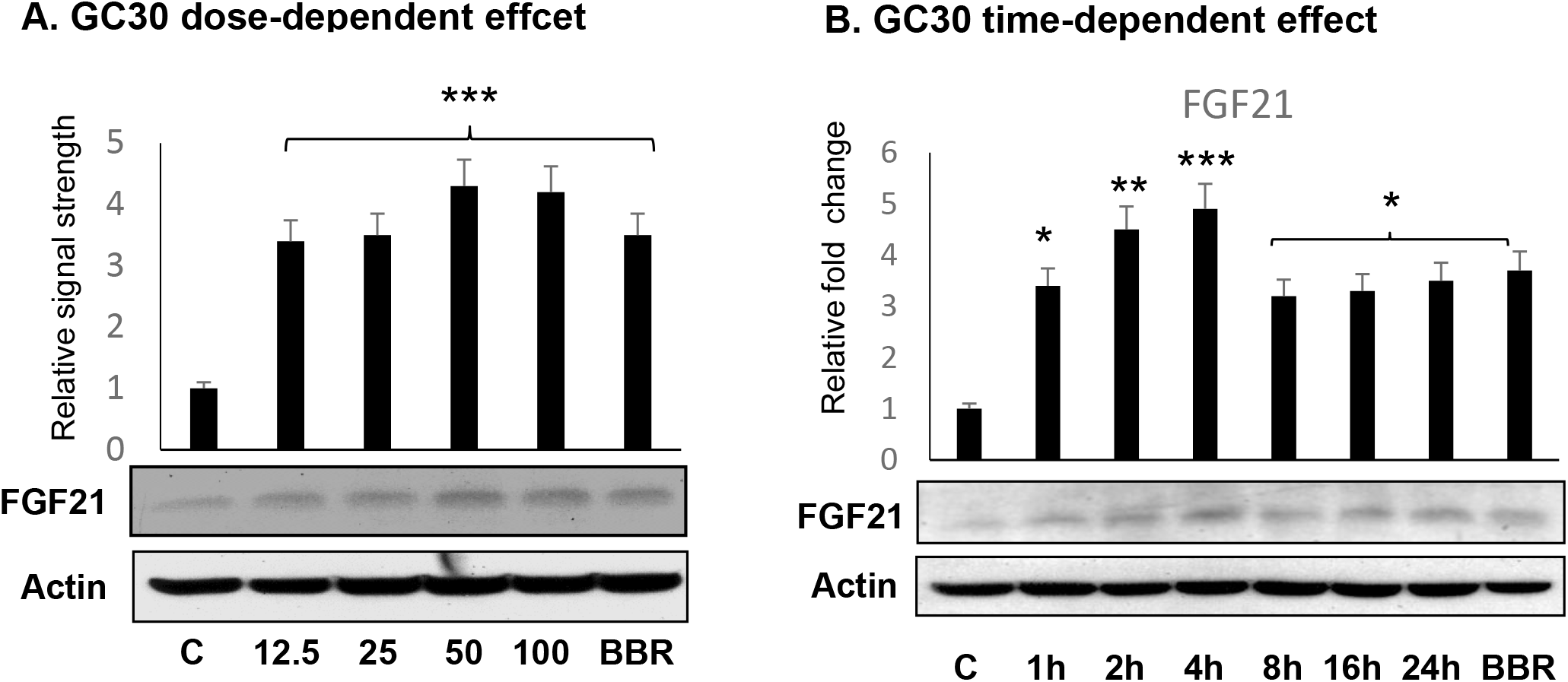
Induction of FGF21 expression in hepatocytes by GC30. FGF21 expression was examined in 1c1c7 hepatocytes after GC30 treatment. A. FGF21 protein expression. The whole cell lysate was used in the Western blot for FGF21 protein. The dose-dependent study was done with GC30 treatment for 4 hr. B. FGF21 protein in a time-course study. GC30 (100 μg/ml) was used to treat the cells for 24 hr. The bar figure represents mean ± SE (n=3). ^*^ p < 0.05; ^**^ p < 0.01; ^***^ p <over the control.

### Induction of FGF21 gene promoter by GC30

FGF21 expression is regulated at the transcriptional level. mRNA and the gene promoter were examined to understand the mechanism of GC30 activity. An increase in mRNA was observed in FGF21 in response to GC30 treatment in dose- and time-dependent manners (Fig. 4, A and B). The gene promoter activity was examined using a luciferase reporter controlled by the FGF21 promoter DNA ^[15]^. The reporter activity was induced by GC30 with 200% elevation over the control (Fig. 4C). In the positive control, the reporter activity was induced by Forskolin through activation of cAMP/PKA pathway (Fig. 4D). The GC30 activity was examined in the regulation of cAMP/PKA pathway with phosphorylation of the transcription factor CREB. An increase was observed in the phosphorylation in GC30-treated cells (Fig. 4E). The GC30 activity was similar to that of Forskolin (Fig. 4E). These results suggest that GC30 may induce FGF21 transcription in hepatocytes through activation of the cAMP/PKA signaling pathway.

**Fig 4.**
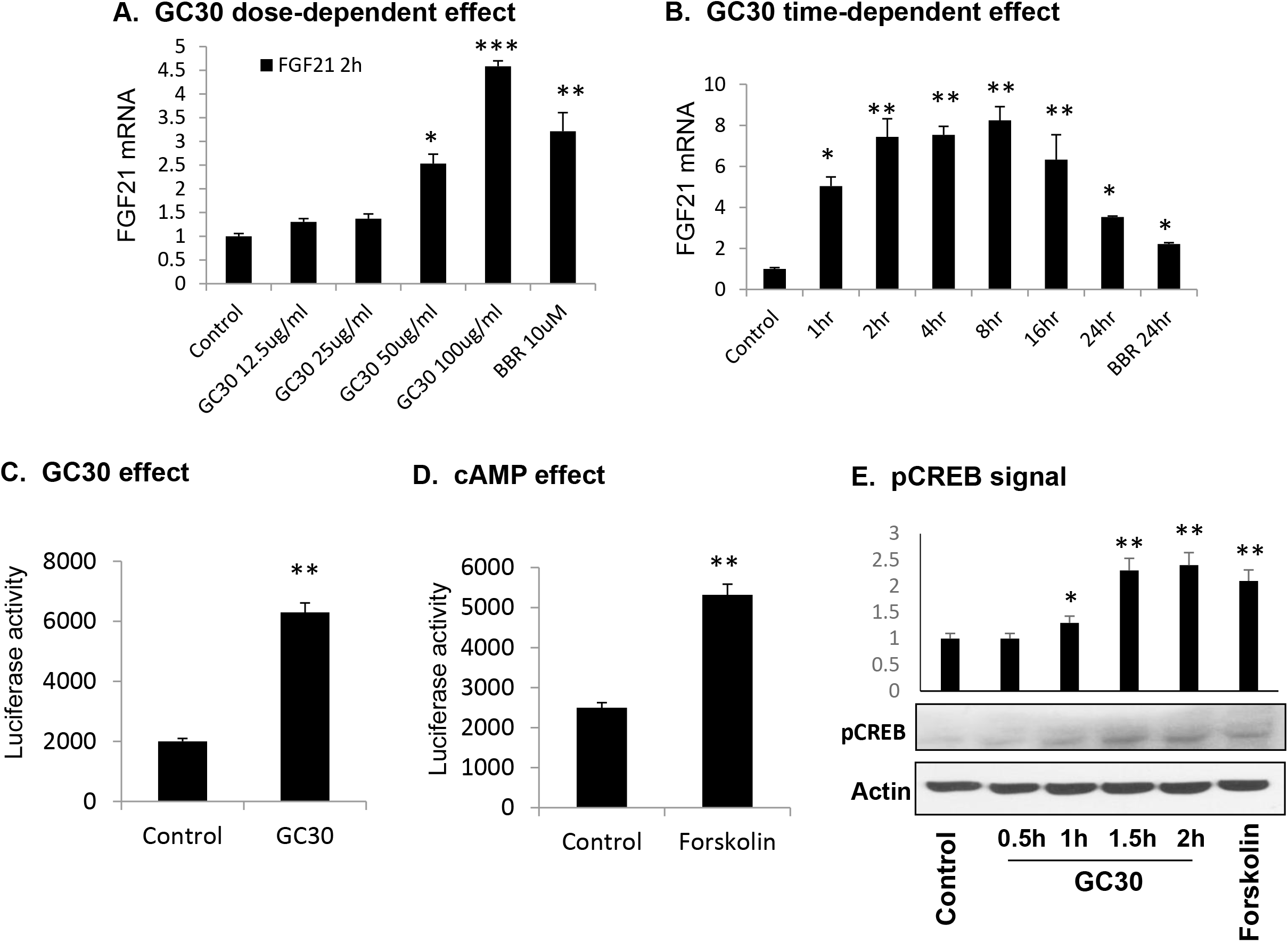
Activation of FGF21 gene transcription by GC30. A. Induction of FGF21 mRNA by GC30 in the dose-dependent study. The FGF21 mRNA was determined in 1c1c7 cells following a 2 hr treatment with different dosages of GC30. mRNA was determined using qRT-PCR. B. Induction of FGF21 mRNA by GC30 in the time-dependent study. The FGF21 mRNA was determined in 1c1c7 cells with GC30 (100 μg/ml) treatment over 24 hr. C. Induction of the FGF21 luciferase reporter by GC30. The FGF21 reporter was transiently transfected into 293 cells, and the cells were treated with GC30 (50 μg/ml) for 16 hr. D. Induction of the FGF21 gene promoter activity by Forskolin. The reporter activity was determined after Forskolin (10 μM) treatment of the transfected cells. E. Induction of CREB phosphorylation by GC30. The phosphorylation was determined in whole cell lysate by Western blotting after treatment of the cells with GC30 (50 μg/ml) and Forskolin (10 μM). In the bar figure, the results are mean ± SE (n=3). ^*^ p < 0.05; ^**^ p < 0.01; ^***^ p < 0.001 over the control.

### Activation of AMPK and inhibition of ATP by GC30

AMPK is an energy sensor in cell whose activity is induced by a reduction of the intracellular ATP level. AMPK activity was examined to understand the mechanism of GC30 activity in the inhibition of hepatic steatosis. GC30 activity was examined in the regulation of AMPK activity by the phosphorylation status (pAMPK) in the cells. The pAMPK signal was induced in the cells treated by GC30 (Fig. 5A). BBR and metformin were used as the positive controls in the activation of AMPK (Fig. 5A). The GC30 activity in the activation of AMPK was not as strong as those of BBR or Metformin. In the study, GC30 was compared with another extract of Kursi Wufarikun Ziyabit (GC70), and other compounds including GN (Geraniin) and EA (Ellagic acid). The pAMPK signal was induced by all of them (Fig. 5A). The data suggest that GC30 activated AMPK in the hepatocytes.

**Fig 5.**
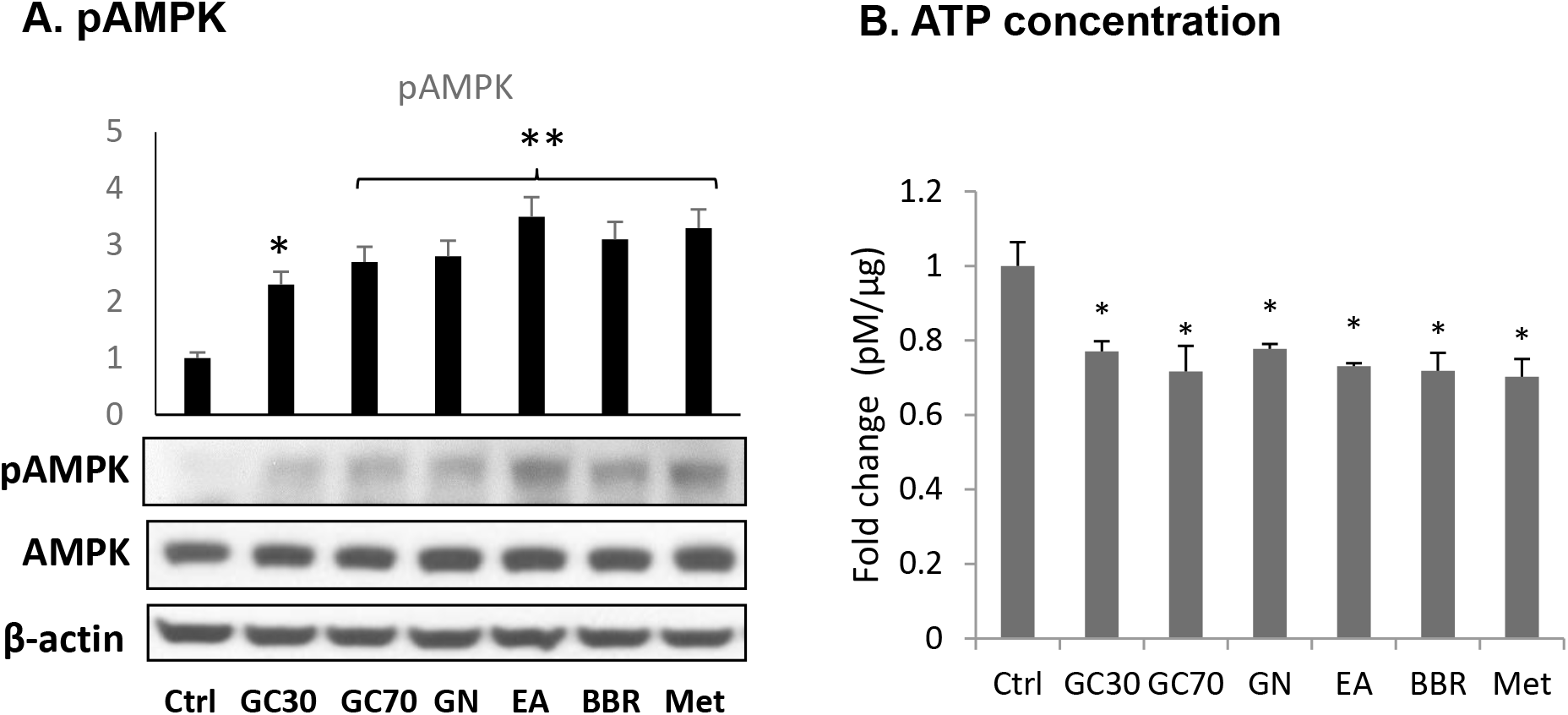
Induction of AMPK phosphorylation and inhibition of ATP by GC30. A. Induction of AMPK phosphorylation by GC30. The HepG2 cells were treated with GC30. The phosphorylation of AMPK was examined in the whole cell lysate at 60 mins after exposure to GC30 (50 μg/ml), GC70 (50 μg/ml), GN (10 μM), and EA (10 μM) by Western blotting. BBR (10 μM) and Met (10 μM) was in the positive controls as AMPK activators. B. Inhibition of ATP by GC30. ATP was determined in the whole cell lysate after GC30 treatment of HepG2 cells. The fold change of ATP is presented in the unit of μM/g protein. In the bar figure, the results are mean ± SE (n=3). ^*^ p < 0.05; ^**^ p < 0.01 over the control.

Intracellular ATP was determined in the cells to understand the mechanism of AMPK activation by GC30. The ATP content was decrease by GC30, which was similar to those of other compounds including GC70, GN, EA, BBR and Metformin (Fig. 5B). These data suggest that GC30 may activate AMPK through reduction of ATP level in the cells.

### Inhibition of mitochondrial ATP production by GC30

Mitochondria are the major source of ATP in hepatocytes. Mitochondrial function was examined to understand the ATP reduction in hepatocytes by GC30. The oxygen consumption rate (OCR) is an indicter of mitochondrial function as the mitochondrial activity is dependent on oxygen. OCR was measured in the GC30-treated cells and a significant reduction was observed in the basal and maximal (uncoupling) conditions (Fig. 6A). The ATP production is catalyzed by ATP synthase in mitochondria. Inhibition of the enzyme by oligomycin leads to a decrease in OCR. The difference of OCR at the basal and oligomycin-treated conditions represents the ATP production capacity in mitochondria. Oxygen is also used in the heat production (uncoupling) by mitochondria. In response to the chemical uncoupler FCCP, OCR is induced for the maximal capacity of oxygen consumption in mitochondria. The difference between the basal and maximal OCR represents the spare capacity of mitochondria. ATP production and spare capacity were all decreased in the hepatocytes by GC30 (Fig. 6B). In the study, BBR was used as the positive control. The GC30 activity was similar to that of BBR in this assay (Fig. 6B). These data suggest that GC30 inhibited ATP production by mitochondria in the hepatocytes.

**Fig 6.**
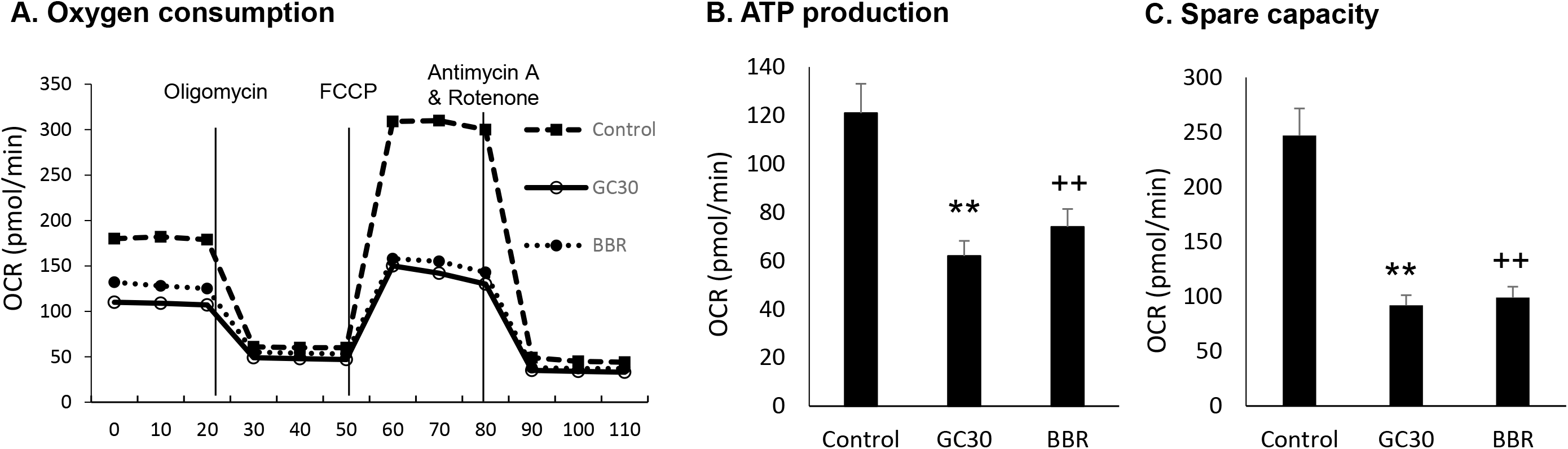
Inhibition of mitochondrial ATP production by GC30. A. Dynamic change of oxygen consumption rate (OCR) of mitochondria in response to GC30. OCR was determined in the HepG2 cells following GC30 (50 μg/ml) for 1 hr. In the positive control, BBR (10 μM) was used. OCR was determined using the XF24 Analyzer over 110 minutes. B. ATP production. The value was a result of OCR difference before and after oligomycin treatment. C. Spare capacity of mitochondria. This was a result of OCR difference between the basal and maximal conditions. The results are mean ± SE (n=3). ^**^ p < 0.01; ^++^ p< 0.01 relative to the control.

### Compound profile of GC30

GC30 was analyzed with liquid chromatograph-mass spectrometry to determine the compound profile. Twenty compounds were identified in GC30 according to the major peaks in the fingerprints of GC30. Those include Quercetin, Quercetin derivative and others. Quercetin provides a basis for certain activities of GC30 in the regulation of ATP production.

## Discussion & Conclusions

Our data demonstrates that the GC30 extract of herbal medicine of Kursi Wufarikun Ziyabit is able to inhibit hepatic steatosis and induce FGF21 expression in hepatocytes. The medicine has been used for hundreds of years in the treatment of diabetes and liver diseases in forks medicines of the countries of Central Asia. However, the cellular and molecular mechanisms were not known for the therapeutic activities. To address this issue, we prepared the GC30 extract of Kursi Wufarikun Ziyabit, and tested its activity in the hepatocyte model. GC30 was found to have activities in the inhibition of development of hepatic steatosis and induction of FGF21 expression in the hepatocyte model. The data suggest that liver is a potential target of the herbal medicine in the treatment of diabetes and liver disease. The mechanism is related to improvement of hepatocyte function by inhibition of steatosis and induction of the endocrine activity. These observations provide a cellular mechanism for the therapeutic activities of the herbal medicine.

Our data suggests that in hepatocytes, GC30 activated the AMPK and cAMP/PKA signaling pathways to generate the pharmacological activities. The AMPK pathway provides a mechanism for inhibition of hepatic steatosis. In cells, AMPK is activated by lack of fuel supply in the stress conditions. AMPK activation acts to restore the fuel supply in cells through an impact in multiple metabolic pathways ^[2]^. Inhibition of the de novo lipogenic pathway by AMPK shifts energy metabolism from anabolism to catabolism, which includes induction of utilization of glucose and fatty acids in the production of ATP. AMPK activation by GC30 represents a molecular mechanism for the GC30 activity in the inhibition of hepatic steatosis. The cAMP/PKA pathway stimulates fuel mobilization and utilization in ATP production through stimulation of glycose and fatty acid oxidation in mitochondria. It also induces FGF21 expression in hepatocytes. Those activities support another molecular mechanism of the GC30 activity in the inhibition of hepatic steatosis. Although the cAMP/PKA pathway induces gluconeogenesis in hepatocytes, the activity may be offset by the activation of AMPK in the GC30-treated cells. AMPK is known to inhibit gluconeogenesis in the liver. Activation of the two pathways (AMPK and cAMP/PKA) provides two molecular mechanisms for the activities of GC30 in the regulation of hepatocyte metabolism.

GC30 exhibited an activity in the suppression of ATP production in mitochondria, which was similar to those of BBR and metformin. BBR and metformin are widely used in the control of blood glucose in the diabetic patients worldwide. They share an activity in the activation of the AMPK signaling pathway by inhibition of ATP production in mitochondria ^[17]^. In current study, GC30 was found to reduce the intracellular level of ATP by suppression of mitochondrial oxygen consumption, which provides a mechanism for the activation of AMPK and cAMP/PKA pathways. Activation of the two pathways is likely a compensatory response to the ATP depletion in favor of ATP production. A similar reaction was reported in our previous study of BBR, which inhibited ATP production by mitochondria with a reduction in oxygen consumption ^[18]^. Interestingly, such an activity was also reported for metformin by other group ^[19]^. Consistently, ATP enhances de novo lipogenesis and gluconeogenesis in liver in the pathogenesis of insulin resistance ^[20, 21]^. Inhibition of ATP production by an uncoupler (2,4 dinitrophenol, DNP) prevented insulin resistance in dietary obese rats ^[21]^. Elevation of intracellular ATP promotes insulin resistance and inflammation in previous studies by us ^[14, 22]^. Suppression of ATP production by mitochondria provides a cellular basis for the GC30 activities in the activation of AMPK in hepatocytes. The molecular target of GC30 remains to be identified in mitochondria.

The compound profile was examined in GC30 using the liquid chromatography-mass spectrometry (LC-MS) technology. Twenty compounds were identified in GC30, in which quercetin and its derivatives were found. Quercetin has an activity in the inhibition of mitochondrial function to decrease ATP production ^[23]^, which may contribute to the GC30 activity in the regulation of AMPK. Quercetin inhibits lipogenesis in adipocytes ^[24]^, which is consistent with the GC30 activity in the inhibition of lipogenesis in hepatocytes.

In summary, we investigated the activity of herbal medicine of Kursi Wufarikun Ziyabit in the regulation of hepatocyte metabolism in this study. The study was conducted with the GC30 extract in hepatocytes. GC30 inhibited the development of hepatic steatosis and induced expression of FGF21 through a transcriptional mechanism in hepatocytes. The activities were associated with activation of the AMPK and cAMP/PKA pathways. GC30 inhibited ATP production in mitochondria to activate the two signaling pathways through ATP depletion. The GC30 activities are similar to those of BBR and metformin. Quercetin was identified in GC30 together with other 19 compounds to support the mechanism of GC30 activities. The study suggests that the herbal medicine may generate the therapeutic activities by targeting liver. However, this possibility remains to be tested in vivo.

## Acknowledgments

This work was supported by the West Light Foundation of the Chinese Academy of Sciences (Grant No. 2016-QNXZ-B-4, 2018-YDYLTD-001) and the Central Asia Drug Research and Development Center of Chinese Academy of Sciences.

## Author contribution

MY, LJ, XX, MS and YN conducted the experiments and analyzed the data. JY and HAA had the concept of the study, designed the experiments. JY wrote the manuscript. HAA is the guarantor of this work and has full access to all the data in the study and takes responsibility for the integrity of the data and accuracy of the data analysis. All authors reviewed the results and approved the final version of the manuscript. There is NO conflict of interest to disclose.

## Notes

**Conflict of interest**: There is **NO** conflict of interest to disclose. All authors have read the manuscript and reached an agreement in publication of the study. The manuscript has not been submitted to other journals.

**Financial support**: This work was supported by the West Light Foundation of the Chinese Academy of Sciences (Grant No. 2016-QNXZ-B-4, 2018-YDYLTD-001) and the Central Asia Drug Research and Development Center of Chinese Academy of Sciences.

### Competing Interest Statement

The authors have declared no competing interest.

